# Polymer Collapse & Liquid-Liquid Phase-Separation are Coupled in a Generalized Prewetting Transition

**DOI:** 10.1101/2024.04.29.591767

**Authors:** Mason N. Rouches, Benjamin B. Machta

## Abstract

The three-dimensional organization of chromatin is thought to play an important role in controlling gene expression. Specificity in expression is achieved through the interaction of transcription factors and other nuclear proteins with particular sequences of DNA. At unphysiological concentrations many of these nuclear proteins can phase-separate in the absence of DNA, and it has been hypothesized that, in vivo, the thermodynamic forces driving these phases help determine chromosomal organization. However it is unclear how DNA, itself a long polymer subject to configurational transitions, interacts with three-dimensional protein phases. Here we show that a long compressible polymer can be coupled to interacting protein mixtures, leading to a *generalized prewetting* transition where polymer collapse is coincident with a locally stabilized liquid droplet. We use lattice Monte-Carlo simulations and a mean-field theory to show that these phases can be stable even in regimes where both polymer collapse and coexisting liquid phases are unstable in isolation, and that these new transitions can be either abrupt or continuous. For polymers with internal linear structure we further show that changes in the concentration of bulk components can lead to changes in three-dimensional polymer structure. In the nucleus there are many distinct proteins that interact with many different regions of chromatin, potentially giving rise to many different Prewet phases. The simple systems we consider here highlight chromatin’s role as a lower-dimensional surface whose interactions with proteins are required for these novel phases.

## INTRODUCTION

Long biopolymers, DNA & RNA, encode genetic information in their sequence. The three dimensional structure and arrangement of these polymers plays an important role in cellular processes. In the nucleus of eukaryotes, the chromosome displays three dimensional structure at many distinct scales, from TADs to euchromatin/heterochromatin [1], and in some cases the colocalization of enhancers, promoters, and otherwise distal genetic elements has been shown to be a key step in initiating transcription [2–4]. While some of this structure is driven by enzymatic activity from loop extrusion factors (LEFS), helicases, isomerases, and polymerases [5, 6], some is likely to be thermodynamic in origin, driven by energetic interactions between long polymers and proteins, small RNAs and other macromolecules [7, 8].

Many nuclear components including transcription factors and heterochromatin-associated proteins have a tendency to phase separate into coexisting liquid phases when isolated at high concentrations, even in the absence of DNA [7, 9, 10]. This phase separation is likely driven by large numbers of weak but specific interactions between these components, which are sufficiently strong to overcome the entropic cost of phase separating, but weak enough to allow for mobility of individual components. Phase separation of protein components is a driving force behind a range of functional structures in the cytoplasm, and has been argued to play a similar role in organizing chromatin [2, 11–1

But there are other thermodynamic drivers of spatial organization available to long polymers like chromosomes and RNA. In particular, long polymers can undergo thermodynamic transitions of their own. Collapse transitions separate a regime where typical configurations are extended from where polymers are condensed and space filling [14]. These transitions can be accessed by solvent quality; in good solvents polymers typically have a weak contact repulsion and exist in an Extended configurational phase. In poor solvents, polymers prefer to minimize their solvent interaction by maximizing self-contacts, leading to Collapsed phases. In the nucleus more complex interactions could in principle drive more interesting configurational phases.

We recently argued that a range of structures at the plasma membrane are *Prewet* : surface phases formed by a combination of membrane mediated forces and interactions between cytoplasmic proteins [15]. In classical prewetting, a thin film resembling a three dimensional phase can adhere to a surface outside of a thermodynamic regime where that bulk phase is stable [16, 17]. But in the biological case, the solid surface is replaced by the fluid plasma membrane, itself found near a 2D liquid-liquid demixing critical point [18]. We showed that in this system the prewetting transition of the bulk merges with the de-mixing transition at the surface, allowing for phase transitions that would not occur if the membrane or the bulk were isolated from each other. This Prewet phase can also be viewed from the perspective of the membrane with bulk proteins shifting the location of the two dimensional liquid demixing transition.

Here we explore a model in which bulk proteins prone to phase separation play an analogous role modulating the properties of a polymer collapse transition. In this *generalized prewetting* transition we consider a long polymer in a good solvent that interacts with bulk components. As we will show, this system has a phase in which bulk components are at near condensed density, and in which the polymer is Collapsed, at parameters in which the polymer would remain Extended in the absence of bulk, and the bulk would remain well-mixed in the absence of a polymer to scaffold it. We investigate the phase diagram of this generalized prewetting transition with both a lattice model and a mean field theory, and explore its implications for genomic structure.

## RESULTS

### Monte-Carlo Simulations

#### Model

We model a single self-avoiding polymer on a 3-D cubic lattice of linear dimension L = 64. A spin variable at lattice site i, 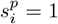 for sites which the polymers passes through and is 0 otherwise. The energy of configurations of this system is given by the Hamiltonian H_poly_:

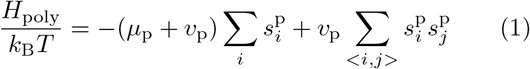

Here both sums are over lattice sites, and < i, j > represents a sum over nearest neighbors in space. µ_*p*_ is a monomer chemical potential which multiplies 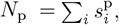, the total number of monomers in the polymer. *v*_p_ is a monomer-monomer interaction energy, and the subtraction in the first term avoids counting contacts between sequential monomers. All simulations take place on periodic lattices, and are constrained to polymer configurations with initial position (*L*/2, *L*/2, 0) and final position (*L*/2, *L*/2, *L*). We also enforce that the polymer have length shorter than *N*_max_, typically 1500 monomer units, so that the polymer obeys *L* ≤ *N*_p_ < *N*_max_. In most of our simulations *N*_p_ ≈ *N*_max_, so that we effectively sample a fixed length distribution, but we allow small fluctuations for numerical convenience.

We couple this single long polymer (yellow in Figure 1A) to a solution of two types of shorter polymers which we term *bulk molecules* (red and blue in Figure 1A) and which are held at fixed chemical potential *µ*_b_ and fixed length *N*_b_. Following our previous work [15], we model the bulk as a mixture of two types of molecules prone to condensing together. Molecules of the same type are self-avoiding while molecules of different types may occupy the same site, and have an on-site attractive interaction with energy *J*_bulk_. Their occupancies at site i are defined analogously to the long polymer by two additional spin variables 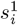 and 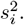. All molecules interact with a weak nearest-neighbor energy *J*_nn_. The bulk Hamiltonian thus reads:

**FIG. 1.**
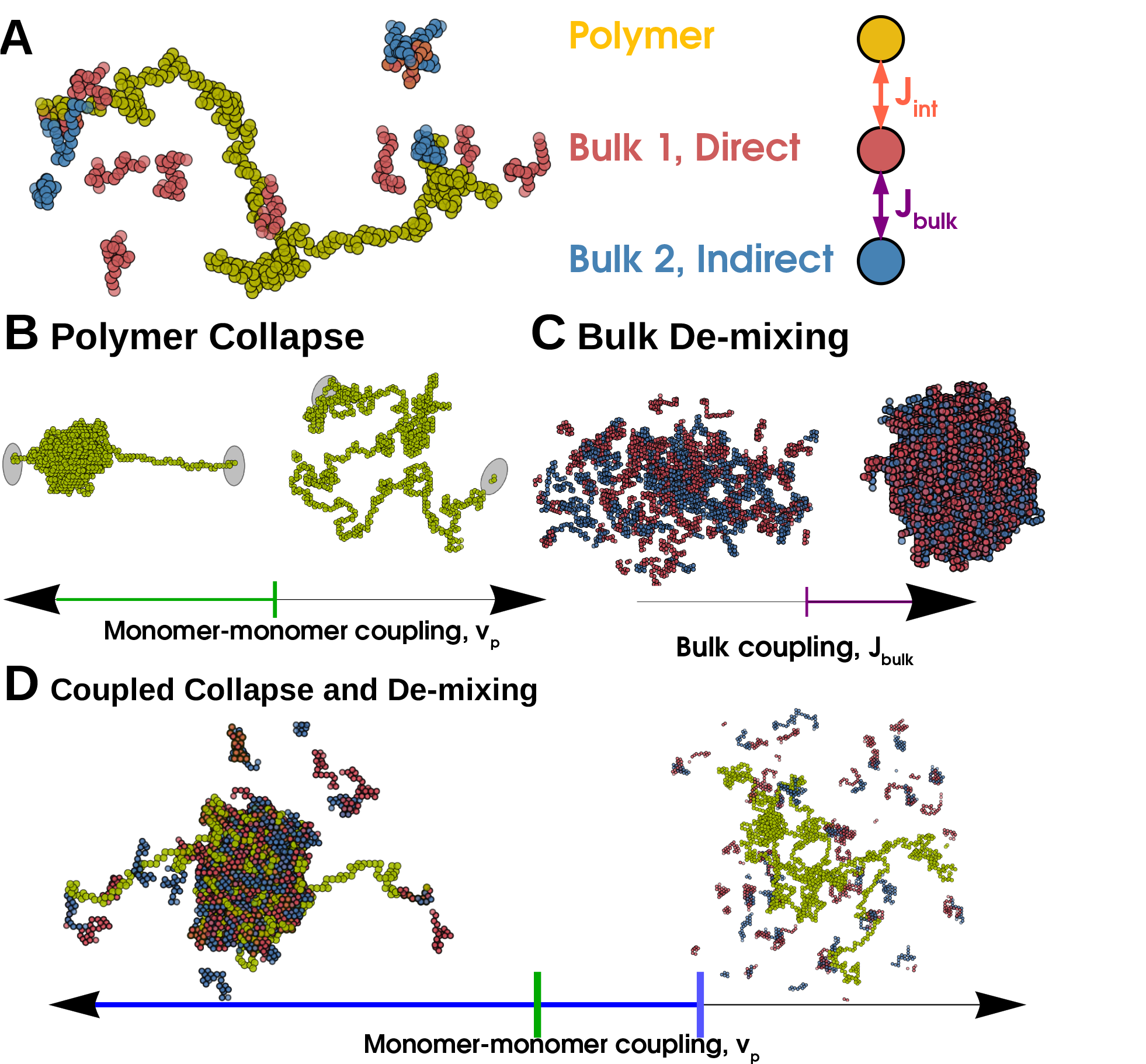
Polymer collapse and phase-separation are coupled in a Generalized Prewetting phase transition. (A) Model coupling polymer collapse and liquid-liquid phase-separation. The long polymer, yellow, interacts with the bulk, red and blue molecules. The polymer is self-avoiding but its monomers have nearest-neighbor interactions that can be attractive or repulsive; the red and blue bulk molecules interact with each other with energy *J*_bulk_. The red molecule and the long polymer interact with energy *J*_int_ and the blue molecule does not interact with the long polymer. (B) A long polymer undergoes a Collapse transition where the polymer’s configuration changes rapidly but continuously when monomer-monomer coupling changes sign, green line. (C) The 3D bulk de-mixes through a first-order phase transition driven by interaction (or composition). (D) When polymer and bulk systems interact, the collapse transition and bulk demixing coincide in a Generalized Prewetting transition, illustrated below as the blue line, which occurs at monomer-monomer couplings more repulsive than those required for collapse in the absence of bulk molecules. The corresponding uncoupled system would have a single Dilute bulk phase and an Extended polymer.

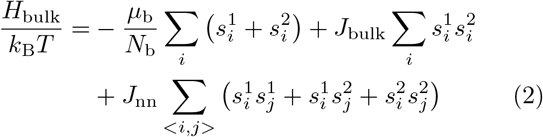

The bulk system corresponds to proteins such as Transcription Factors that interact with each other and with chromosomes. *J*_bulk_ represents the interaction between these proteins, modulated by post-translational modifications such as phosphorylation. And a chemical potential *µ*_b_ sets the concentration of these proteins.

The long polymer and one of the bulk molecules, red in the Figures, are coupled by interactions of strength *J*_int_, giving an interaction energy:

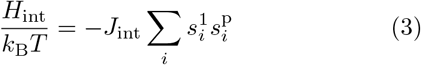

*J*_int_ can be interpreted as the affinity of a protein for a specific DNA sequence or histone modification. With these interactions, the red bulk molecule could correspond to a transcription factor and the blue to a co-activator such as Mediator. We illustrate this model in Figure 1A.

We sample equilibrium configurations of this system using a Monte-Carlo procedure, fully explained in the Methods. Briefly, we generate a long polymer through a set of three elementary moves corresponding to the addition, deletion, and ‘kink’ of a given bond. Bulk molecules are updated with reptation and kink moves that conserve length, and we exchange these particles with a reservoir. We accept moves with Metropolis probability chosen to obey detailed balance given the above Hamiltonian.

#### Thermodynamics of an Isolated Polymer

In the absence of the bulk system, the long polymer can access several thermodynamic phases. These phases correspond to distributions over the spatial configurations of the long polymer, 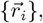, where 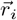 is the posi-tion of the *i*^th^ monomer. For idealized polymers in regions where boundary conditions can be neglected, these phases can be characterized by scaling relationships between *N*_p_ and the radius of gyration *R*_g_, defined by 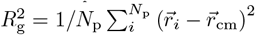 with 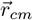 the center of mass of the chain. The relation 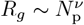 defines the exponent *ν* which characterizes different phases. We show sample configurations of these phases in Figure 1B

When *µ*_*p*_ is negative and *v*_p_ is positive or small and negative, the polymer is in the *Short* phase. Here *N*_p_ is small, ≈ *L*, and the scaling exponent *ν* ≈ 1.; the polymer is a fluctuating chain of length *N* whose end-to-end distance *R*_g_ ∼ *N*_p_. In our simulations we typically set *µ*_p_ = 0, and are not concerned with this *Short* phase.

When *v*_p_ is large and attractive, the polymer is in the *Collapsed* phase. Here *N*_p_ fluctuates about *N*_max_, and the scaling exponent *ν* approaches the space-filling limit of 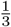 in 3D. We show a typical Collapsed phase in 1B.

When *v*_p_ is large and repulsive, but *µ*_*p*_ is positive, the polymer is in the *Extended* phase. Here the length fluctuates about *N*_max_, but the polymer configuration is Extended and the average density of the polymer approaches 0 as *N*_max_ approaches infinity. The Extended and Collapsed phase are separated by a line of continuous transitions at *v*_p_ = 0. We show a typical Extended configuration in 1B.

#### Bulk Molecules Phase Separate and Modify the Polymer Phase Diagram

Our bulk system has phases and phase-transitions independent of its interactions with the long polymer. The bulk phases are *dilute* and *dense* corresponding to gas and liquid-like states. At certain chemical potentials, or densities, these bulk phases will coexist and the system phase-separates, drawn as the yellow region in Figure 2E, with typical configurations in each phase shown in Figure 1C.

**FIG. 2.**
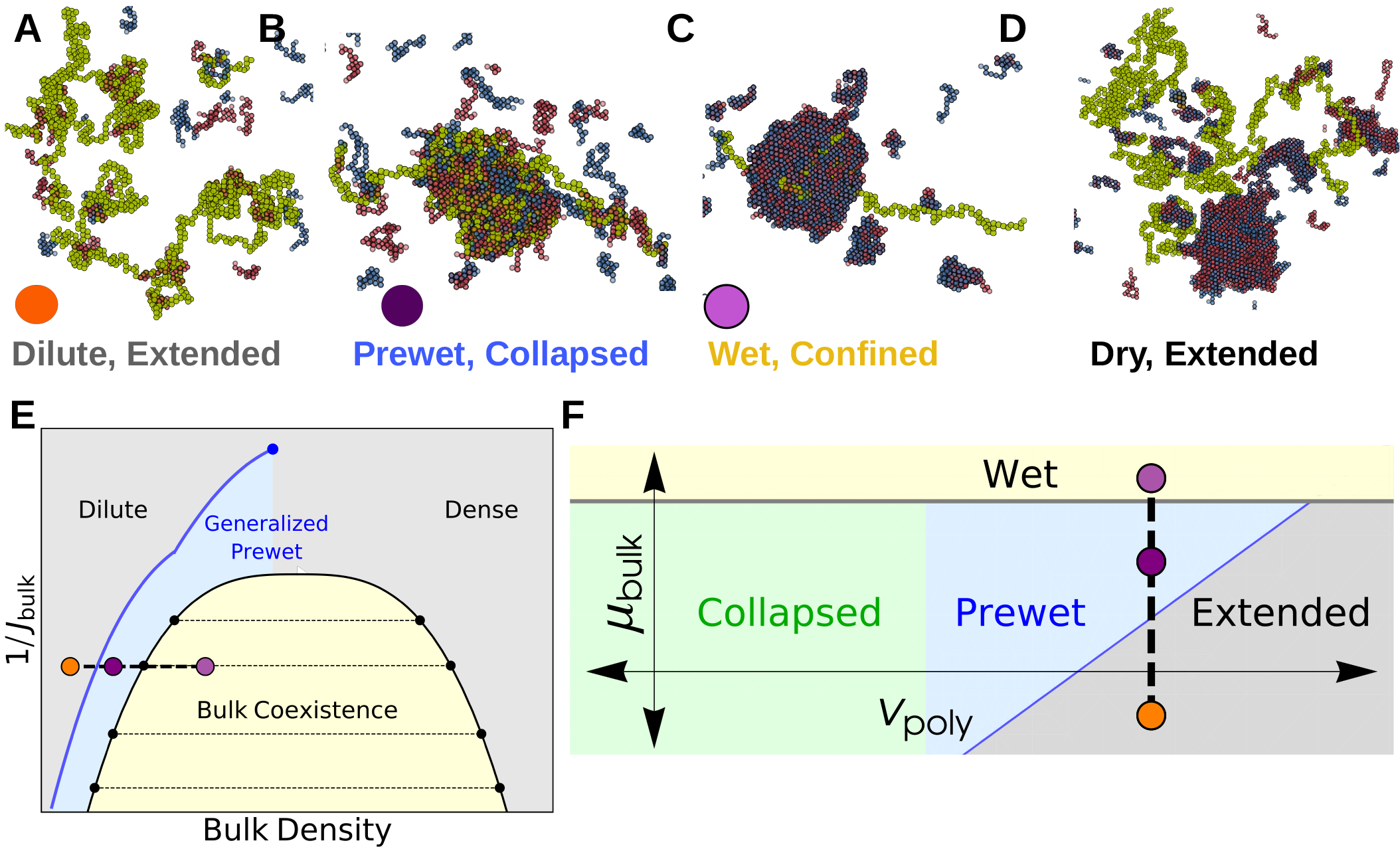
The bulk system shifts the Polymer Collapse phase boundary. (A-D) Configurations of coupled phases: Outside of the bulk coexistence region there are two phases that correspond to phases of an isolated polymer: (A) Extended and (B) Collapsed. The Collapsed phase coincides with condensation, or Prewetting, of the bulk fluid. The Extended phase has corresponding Dilute bulk phase. Within the bulk coexistence region the polymer may prefer to enter the droplet, confining itself (Wet, C) or avoiding it (Dry, D). (E) Schematic bulk phase diagram: Generalized Prewet phases occur outside of the bulk coexistence region and terminate in a critical point above the bulk critical temperature. Colored points correspond to above configurations. (F) Schematic phase diagram of monomer-monomer interactions *v*_p_ and bulk chemical potential *µ*_b_. The bulk modifies the phase boundary of the polymer, allowing it to collapse in regions where it otherwise would not; this Collapse transition coincides with condensation of the bulk on the surface of the polymer.

Within the bulk coexistence region the polymer can prefer to localize to the dense phase or the dilute phase. This transition occurs when the energy of polymer-bulk interactions surpasses the entropic and energetic penalties of confining the long polymer to the bulk dense phase. This transition is analogous to the Wetting transition: when *J*_int_ is strong the polymer prefers to enter the droplet to maximize its interactions with the bulk dense phase. We show sample configurations from *Dry* and *Wet* phases in Figure 2C,D. The polymer configurations in the Wet phase depend on *J*_int_ and *J*_nn_, which together determine solvent quality of the dense phase. A polymer that prefers to be Extended in an infinite bulk dense phase can be confined by a finite-size droplet separate from the collapse transition.

Outside of the bulk coexistence region we find another phase where the polymer is Collapsed and also enriched with a dense-like phase of bulk molecules, we show a typical configuration in 2B. This is reminiscent of prewetting where a thin bulk domain forms which is adhered to a particular surface phase [15, 17]; we call phases where polymer collapse coincides with polymer-only bulk demixing *Generalized Prewet - Collapsed*. Consistent with this view, the polymer phase-diagram effectively shifts and the collapse transition occurs at repulsive *v*_p_. Polymer collapse and bulk demixing couple because this minimizes the volume of an otherwise unfavorable bulk phase, while maintaining a high density of bulk-bulk and bulk-polymer interactions. Increasing µ_b_, we transition discontinuously between the three phases shown in Figure 2A-C, which we draw schematically in the phase diagrams sketched in Figure 2E,F.

### Mean-Field Theory

Here we develop a mean-field theory that couples a long, collapsible polymer to a bulk fluid with a propensity to phase-separate. We begin by briefly re-deriving the phase-diagram of an isolated polymer chain, then explore the thermodynamics of the coupled system.

#### Isolated polymer

We write the free energy of an isolated polymer chain, 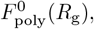, in terms of the order parameter *R*_g_, and at fixed length *N*_p_:

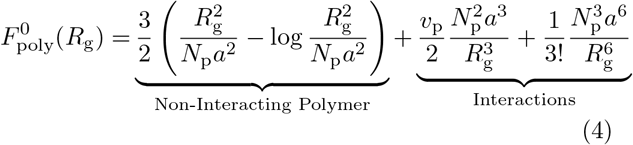

The first two terms labeled ‘Non-Interacting polymer’ are entropic contributions to the free-energy from gaussian statistics, and can be derived by replacing the radius of gyration with the end-to-end distance, with a the linear dimension of a single monomer. A free-energy written in terms of *R*_g_ has additional entropic costs due to confinement – but these, along with higher order interactions, are never relevant so we neglect them here (see Supplement). The final two terms capture mean-field interactions between monomers [19]. The parameter *v*_p_ is the difference between monomer-monomer and monomer-solvent interactions. The point *v*_p_ = 0 separates an Extended polymer phase, where the monomer density 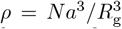 approaches 0, from the Collapsed phase where ρ is finite, see the green line Figure 3A. This transition is continuous but sharp in the large *N*_p_ limit [14, 20].

**FIG. 3.**
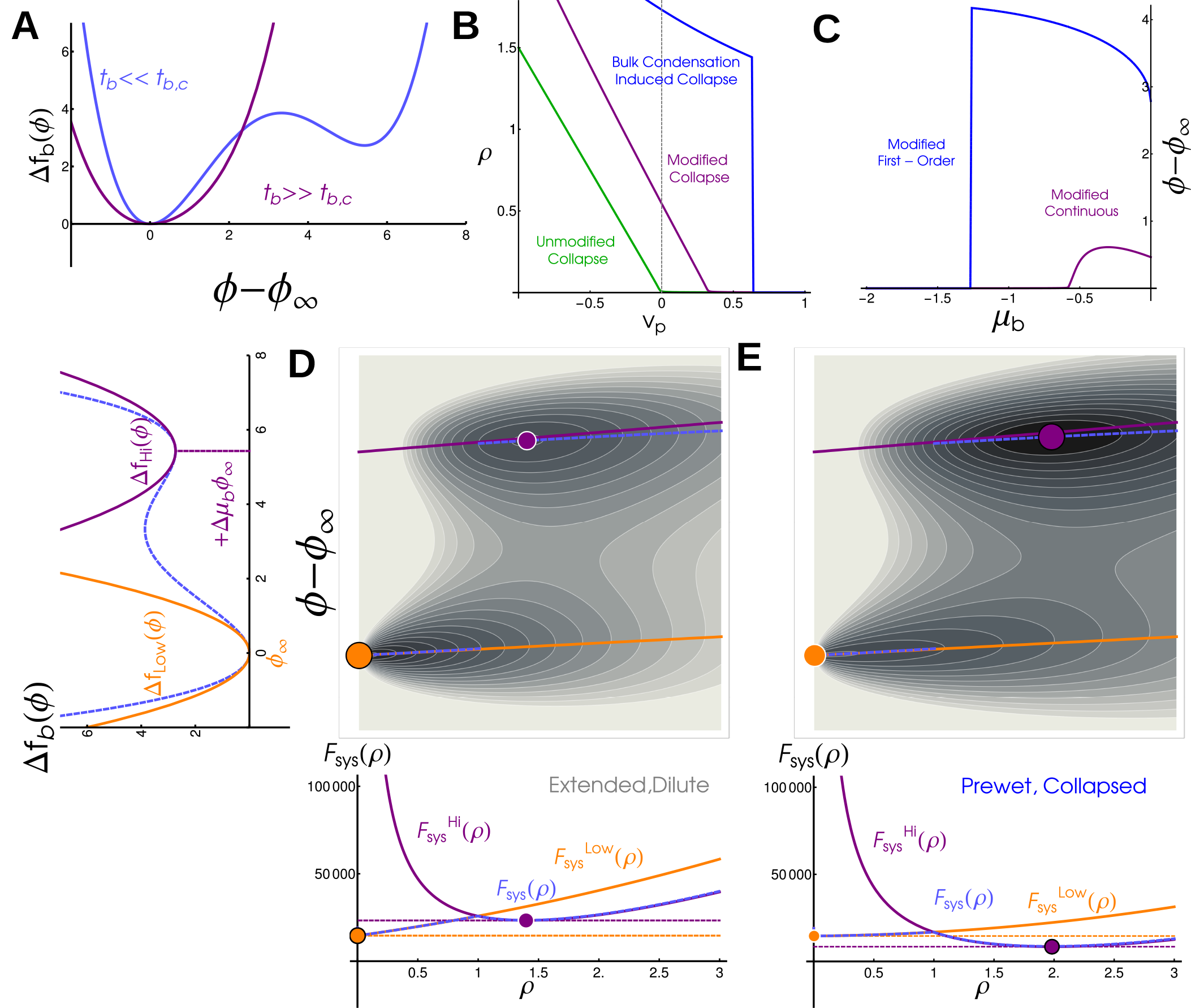
Bulk phase-transitions modify the polymer collapse transition in a coupled mean field theory: (A) Free energy density of the bulk, Δ*f*_bulk_, in super-critical (purple) and sub-critical (blue) regimes. We distinguish these regimes by the presence of one minimum or two well-separated minima. (B) Polymer density *ρ* as a function of monomer-monomer interaction strength *v*_p_. An isolated polymers sees a continuous but sharp collapse transition at *v*_p_ = 0 (green line), but in coupled systems collapse occurs at repulsive *v*_p_ *>* 0 in continuous (purple, bulk has one minima) or discontinuous (blue, bulk has two minima) transitions. (C) Bulk density difference, *ϕ − ϕ*_*∞*_, as a function of bulk chemical potential *µ*_b_. In coupled systems *ϕ − ϕ*_*∞*_ increases with *µ*_b_ in continuous (purple) and discontinuous (blue) transitions. In an uncoupled system bulk condensation occurs at *µ*_b_ = 0 (not visible on this plot). (D) (left) Quadratic approximation to bulk free energy density in the subcritical regime. Orange corresponds to Dilute phase, purple to Dense phase, and dashed blue is the full free energy. (top) Free energy landscape for a system in the Extended, Dilute phase. Local minima corresponding to Extended (orange) and Collapsed (purple) phases are marked. The absolute minima of *ϕ − ϕ*_*∞*_ (dashed, blue) interpolates between the quadratic approximations (solid lines, purple and orange). (bottom) Free energy as a function of polymer density *ρ*. The global minimum of this system is at the low-density, low-*ϕ* minima (orange). E) Energy landscape (above) and free energy as a function of polymer density (below) for a system in the Collapsed, Prewet phase. The global minimum of this system occurs at the high density, high *ϕ* minima.

#### Coupling Polymer to 3D Bulk

We employ a simple mean field theory for an order parameter ϕ which roughly describes the density of bulk phase separating components [21, 22]:

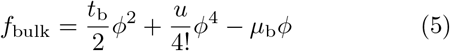

Here *t*_b_ corresponds to the reduced temperature, a measure of interaction strength between bulk molecules, µ_b_ is the chemical potential, and u captures higher order interactions and is required for stability. For *t*_b_ < 0, coexistence occurs at µ_b_ = 0 with energy minima located at 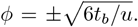. For *t*_b_ ≥ 0 there is one stable phase set by *µ*_b_. These two phases meet in a critical point at *t*_b_ = 0.

In a standard mean field approximation without a polymer, the order parameter is assumed to take the value *ϕ*_*∞*_ which globally minimizes the free energy density f_bulk_. In the presence of a long polymer, we further assume that a sphere of volume 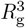 has a polymer density *ρ* and a potentially different value of the order parameter ϕ. Here we assume that both order parameters, *R*_g_ (or equivalently *ρ*) and *ϕ* together minimize the free energy of the coupled system.

We describe interactions between the bulk and the long polymer via an interaction energy density f_int_ and corresponding interaction energy 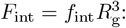:

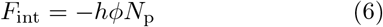

These interactions can be interpreted as the binding energy between the bulk and the long-polymer. To compactly write the coupled free energy of the polymer and its order parameter environment we define Δ*f*_bulk_ = *f*_bulk_(*ϕ*) *− f*_bulk_(*ϕ*_*∞*_). The free energy of the coupled system can then be written as:

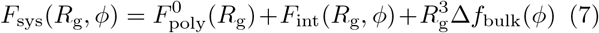

where we have subtracted off a contribution from the polymer-free bulk.

#### A super-critical bulk system modifies the polymer collapse transition

To see how the bulk modifies the thermodynamics of the long polymer, we examine the behavior of a weakly interacting bulk by approximating the free energy as quadratic, 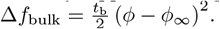. In this quadratic approximation the bulk can be minimized over *ϕ* analytically, leaving just one additional R_g_ dependant contribution to the effective free energy of an isolated polymer, so that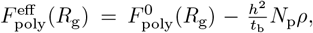, and a constant term *−hϕ*_*∞*_*N*_p_. This additional term has exactly the same form as the monomer-monomer coupling, defining a new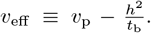. Since there are no other qualitative changes to the free energy, the effect of the bulk in this regime is precisely to shift the parameter regime of the collapse transition, see purple line in Figure 3A. The value of the bulk order parameter tracks with the density of the polymer and sees a sharp, continuous increase through the collapse transition, see Figure 3B, purple line. This effective interaction becomes very strong near the critical point of the bulk, where *t*_b_ is near zero.

#### Strongly interacting bulk drives a first-order transition

In the limit where the bulk system is instead near to phase coexistence, when t_b_ < 0 the bulk free energy density (equation 5) is not well described by a single quadratic minimum. Still, the order parameters *R*_g_ and *ϕ* take values which minimize the combined free energy *F*_sys_ (equation 7). While this minimization cannot be done analytically, minima are easy to find numerically. Contour plots of *F*_sys_ are shown in Figure 3D,E (with *ρ* rather than *R*_g_ on the x axis), where there are two distinct minima. As parameters change, there is an abrupt transition as one minimum becomes lower in free energy than the other. This leads to discontinuities in the polymer density (blue curve in Figure 3B) and in the bulk order parameter (blue curve in Figure 3C).

To gain additional insight into the nature of this abrupt transition, we can also approximate the free energy as quadratic around both dense and dilute phase minima, treating each with the same approximation as above (or-ange and purple lines in Figure 3D,E left). This predicts that local minima will lie near the corresponding orange and purple lines in Figure 3D,E, which meet the y-axis at the locations of the local minima of the bulk free energy density. In this approximation, the system takes the position of either the purple or orange dot, and a first order transition occurs when they have the same value of *F*_sys_. The free energy about the bulk dense phase sees an additional contribution proportional to *R*^3^*µ*_b_*ϕ*_*∞*_. This penalizes Extended configurations and drives the polymer to collapse in an abrupt prewetting transition that accompanies moving to the local minimum associated with the bulk dense phase. Our simulations and theory show that *ρ →* 0 as *µ*_b_ *→* 0. This suggests that, in the thermodynamic limit, the Wet phase corresponds to an Extended polymer localized to the bulk dense phase as long as *h* > 0.

### Multi-component Polymers

In many biological scenarios bulk proteins interact strongly with certain regions of the chromosome, and weakly or not at all with others. We modified our simulations by partitioning the long polymer into discrete segments that either interact with the bulk (yellow in Figure 4A) or do not (green in Figure 4A). We use this to understand how polymer sequence interacts with the behavior we describe above.

**FIG. 4.**
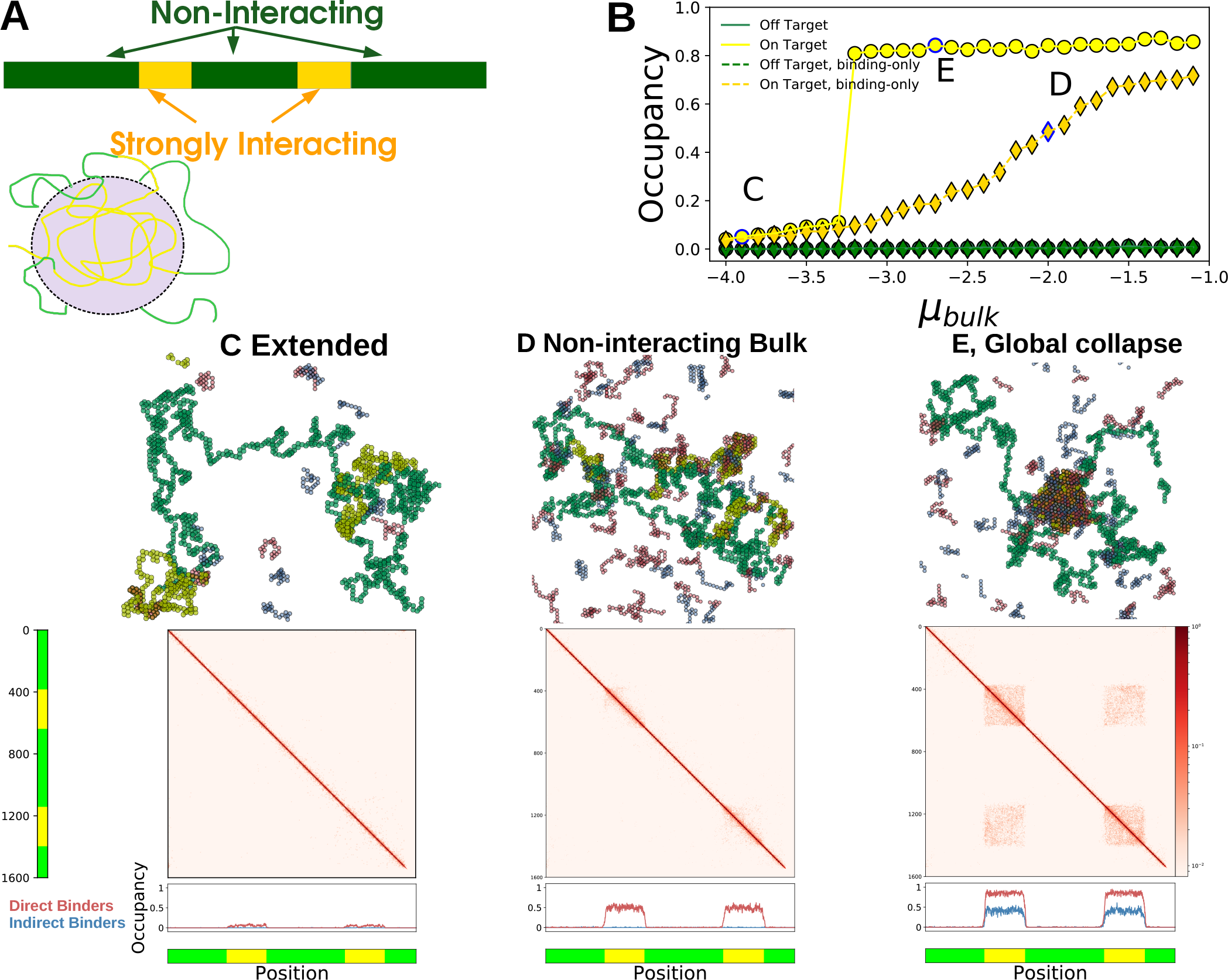
Bulk proteins change the configuration of a multi-component polymer. (A) (upper) Illustration of a multi-component polymer: the polymer has discrete, contiguous segments of monomers that interact differently with the bulk molecules. Here yellow interacts strongly while green is non-interacting. (lower) Schematic showing the selective collapse of yellow segments into a Prewet phase. (B) Occupancy of yellow and green monomers for the bulk binding molecules, as a function of *µ*_b_. Collapse of the yellow segments coincides a jump in the occupancy of yellow segments, but not green. Letters in the plot correspond to the figures below. (C-E) Simulation configurations, monomer contact probabilities, and occupancy profiles of different multi-component polymer phases. (C) Extended phase: the polymer is Extended and bulk occupancy is low and near-uniform. (D) In non-interacting bulk systems, the occupancy of the yellow regions is high (bottom), but the polymer remains in an Extended configuration (middle). (E) Global collapse: (middle) long range, inter-segment contacts are formed via interactions by the bulk phase, red off-diagonal regions in the contact probabilities plot. (lower) Both red and blue bulk molecules are enriched in the yellow segments, but not in the green.

Since the bulk interacts differently with the two monomer types, there are situations where strongly interacting segments collapse but non-interacting segments remain Extended. At low µ_b_ the polymer is Extended (Figure 4C) while at high µ_b_ all interacting segments form a *globally Collapsed* phase (Figure 4E). These phase differ in their occupation of bulk molecules and their polymer-polymer contacts. The Extended phase sees low occupancy of directly binding bulk molecules (red) at the yellow segments (4C, bottom) while both direct and indirect (blue) bulk molecules are enriched in the globally Collapsed phase 4E, bottom). The occupancy of yellow monomers increases discontinuously with µ_b_ while the green monomers remain mostly unoccupied (Figure 4B). When bulk molecules do not interact with each other the occupancy of yellow regions increases smoothly with µ_bulk_ (4B, diamonds), and there is no accompanying occupancy in indirect binders (4D, lower).

Polymer-polymer contacts in the Extended phase are homogeneous, while the globally Collapsed phase sees long-range contacts between monomers of the same type, even those separated by an intervening sequence (4E,middle). In this globally Collapsed phase the reduction in surface tension associated with merging domains must overcome the entropic penalty of looping non-interacting stretches as occurs in our simulations. We also see transient but long-lived locally Collapsed states, and there may be regimes where multiple identical domains stably coexist without merging.

## DISCUSSION

Here we have presented a model where a long polymer near a collapse transition couples to a bulk solvent near a de-mixing transition. We find that this coupling allows both polymer collapse and bulk demixing to occur well outside of their uncoupled phase boundaries. This is reminiscent of prewetting phase-transitions, where a boundary condensed phase is stabilized via interactions with a surface, sometimes undergoing its own transition [15, 17]. Here we discuss the ways biology could use these transitions, and further questions on their physics.

### Transcriptional activation through a prewetting transition

Transcription factors interact directly with enhancers, and many feature ‘activation domains’ that phase-separate with transcriptional machinery such as the co-activator Mediator [7, 23, 24], playing roles roughly analogous to the red and blue polymers in our simulations. These Prewet phases could naturally integrate information from transcription factor concentrations into transcriptional regulation by controlling where along the chromosome Prewet domains occur (Figure 4B). Transcription factor concentration could also modify how enhancers, promoters, and distant genomic regions interact in three dimensions, as in Figure 4E,F. Indeed, many Eukaryotic promoters become spatially proximal to regulatory elements during transcription initiation [2, 3, 25, 26].

### Transcriptional repression with Prewet phases

Many proteins that interact with heterochromatin and other transcriptionally repressed regions have a propensity to phase-separate in the presence or absence of DNA [8, 9, 27–2 Heterochromatin is typically more dense than transcriptionally active chromatin, and is inaccessible to transcriptional machinery [30]. It is possible that heterochromatin is best thought of as a separate generalized Prewet phase, which excludes transcriptional machinery, and is rich in DNA and certain proteins. The complex interactions between the sequence of regulatory elements and cell-type specific transcription factors could regulate transcription by determining which regions are prewet by transcriptionally inaccessible phases [28, 29].

### Prewet phases permit specificity in transcriptional regulation

Eukaryotes face enormous genetic ‘crosstalk’ [31] in that many transcription factors bind promiscuously to many potential genomic targets, so that each individual binding site on the genome carries little specificity. The prewetting phase transition naturally integrates over a long stretch of chromosome, yielding an effective cooperativity in when and where domains appear. While the implications of regulation in this manner is not our focus here, we do expect that long stretches of chromosome with many weak transcription factor binding sites could undergo collapse transitions in a manner that is highly sensitive to small changes in transcription factor concentrations.

### The composition and genomic localization of Prewet domains can encode and read out cellular identity

Cellular identity is thought to be encoded by *core regulatory complexes* (CoRCs) composed of a small number of transcription factors which regulate themselves and downstream cell-type specific genes [32, 33]. Evidence for these complexes comes in part from probes of DNA-Protein interactions. Chromatin Immuno-precipitation (ChIP-seq) studies probe the spatial proximity of specific proteins to specific regions of the genome, and often find long stretches enriched in cell-type specific groups of a few transcription factors. Proteins that map to the same regions are often interpreted as arising from large, macromolecular complexes. But a generalized Prewet phase localized to certain genomic regions has a similar experimental signature, as Figure 4E (occupancy) illustrates. Molecules that do not bind directly to the polymer (blue) localize to the same region of the polymer as molecules that have di-rect interactions (red) so long as both of these molecules phase-separate with each other. Many transcription factors that inspired this model for CoRCs phase-separate at high concentrations (OCT4, NANOG [23]), and are found as large, localized assemblies in some cell types via microscopy. Prewet phases which depend on cellular identity may also explain cell type specific differences in the three-dimensional structure of chromatin (Figure 4E, polymer contacts).

### Phase diagrams for generalized Prewet phases

Past work has investigated how binary bulk mixtures can lead to polymer collapse even when either component on its own does not, a phenomena known as polymer-cononsolvency [34]. As in that work, far above the critical point in our more complicated bulk mixture, the transition from an Extended to a Collapsed state is continuous, though steep. Near the two phase coexistence region of a complicated bulk this transition becomes abrupt, as we see in simulations and mean-field theory. This implies a tri-critical point which we have not thoroughly explored, near but outside of bulk coexistence, where the line of first-order prewetting transitions meets the line of second order transitions. By analogy to prewetting to surfaces, we also anticipate that the line of prewetting transitions meets bulk coexistence at another tricritical point, where Dry, Wet, and Prewet phases coexist. At these tricritical points the polymer likely sees non-trivial scaling behavior. We also expect interesting finite-size effects in the wetting regime when the length of the polymer is such that its Collapsed size is of order the size of the bulk domain. And finally, we anticipate coupling many-component bulk systems to a multi-component polymer will yield very rich physics [35].

### Our model differs from past models of phase sep-aration and chromosomes

Recent work has investigated the role of polymer ‘scaffolds’ on the phase-separation of 3D proteins [10, 36, 37], and how 3D proteins mediate communication between distal genomic regions [38, 39]. This work has emphasized that polymers can widen regimes in which a bulk fluid undergoes a phase-transition. Our results are also related to a body of work investigating how demixing transitions interact with polymer meshes [40, 41], though most of that literature has focused on cytoskeletal meshes, which interact elastically, but do not contribute a transition of their own. Depending on the sign and strength of the interaction between liquid droplets and mesh components, demixing transitions can be enhanced or inhibited, and are generically accompanied by distortions of the polymer mesh. Here we innovate by jointly considering the polymer collapse transition and bulk condensation. Our work is also related to models of the ParABS system for chromosome segregation in bacteria, which explicitly describe the interactions of a dynamic polymer with interacting proteins, but do not focus on the phase behavior of their bulk system [42].

### Distinguishing prewetting from wetting

Although we focus on a region of prewetting phase transitions, our model features other transitions that resemble wetting, where a bulk phase prefers to localize to a surface [16]. Wet phases are stable even without a long polymer; while polymers can partition into them, the presence of a Wet phase does not imply presence of a specific polymer. As such, Wet phases are less suitable for regulation by controlling DNA’s interaction with specific components or its three dimensional structure. Overexpression of transcriptional activators quite generally inhibits transcription through a phenomenon dubbed ‘squelching’ [43]. It is possible that squelching marks a transition into bulk coexistence where macroscopic droplets compete with the DNA for transcriptional machinery.

### Other types of generalized prewetting may be common

This work and a similar model we proposed to explain membrane-localized signaling clusters [15] contain two defining features: (1) a bulk fluid that de-mixes in 3D, and (2) a lower-dimensional surface with a phase transition that is different from that of the bulk. In both of these cases the phase-behavior of the surface gives the bulk qualities of a transition that it didn’t have before – a 2D critical point in the membranes case, and qualities of a polymer collapse transition here. Eukaryotic cells have many surfaces with phase-transitions that do not occur in bulk, and biology often needs to localize processes to specific locations. We speculate that biology uses a variety of these *generalized-prewetting* transitions to accomplish specific localization, and to imbue a bulk fluid with properties such as a diverging susceptibility. Work by ourselves and others [10, 41, 44, 45] will continue to uncover the rich physics allowed by surface phase-transitions.

## Acknowledgements

We thank Asheesh Momi, Yu Fu, Isabella Graf, and Simon Mochrie for feedback on the manuscript, and Pranav Kantroo, Martin Girard, Sarah Veatch, and Simon Mochrie for helpful discussions. This work was supported by NIH R35 GM138341 and NSF 1808551.

## METHODS

### Simulation Model

We sample polymer configurations using a Monte-Carlo procedure [46] composed of three elementary moves that correspond to Addition, Deletion, and Kink of a given bond, where a bond is a link between successive monomers in the polymer, red in Figure 5. Any move may be viewed as the translation of a bond in a direction orthogonal to the direction of the bond (arrows in 5). The type of move proposals available is completely determined by the vertices of two monomers neighboring the monomers participating in the bond. If translation of the bond gives no intersection with neighboring monomers, the move is *Bond Addition* and corresponds to adding two monomers to the chain. If the translated bond intersects with both monomers in the chain the corresponding move is *Bond Removal* and the polymer loses two monomers. If one of the neighboring monomers is intersected by the translated bond, the move is a *Kink* that does not change the length of the polymer, but alters the configuration. These moves are illustrated in Figure 5B.

**FIG. 5.**
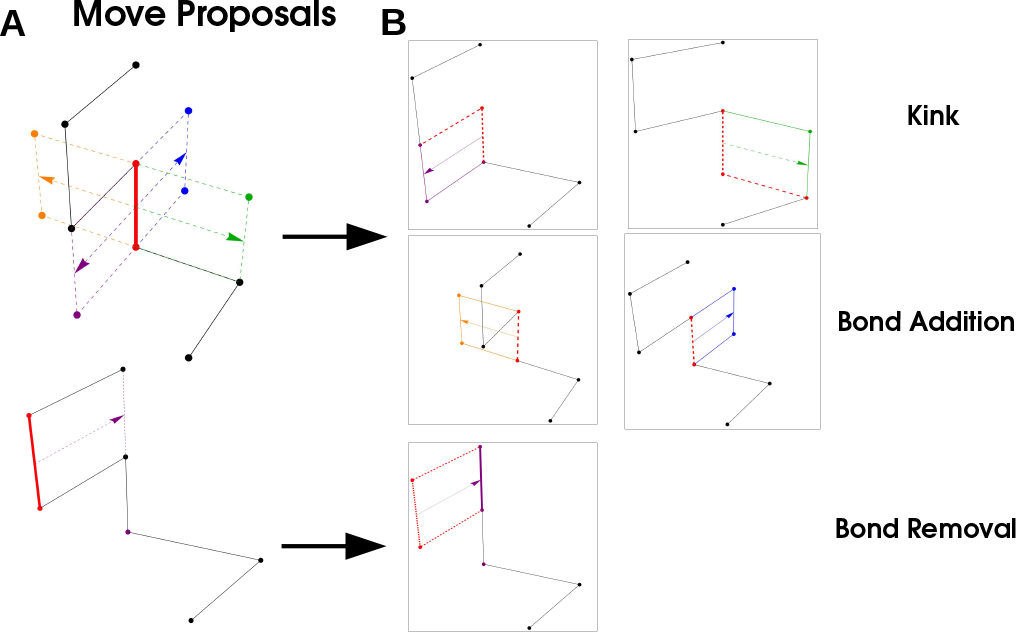
Simulation Procedure: A bond (red) is selected, and translated in a direction orthogonal to its orientation. These moves correspond to a ‘Kink’ (purple and green arrows) where a the polymer configuration changes but the length does not, ‘Bond Addition’ where two monomers are added to the polymer, and ‘Bond Removal’ where two monomers are removed from the polymer

We simulate the polymer as follows: we randomly select a monomer, at position s, and define its bond as the vector between the monomer and the s+ 1 monomer. We then calculate all bond translations and whether they correspond to addition, removal, or kink moves. We then randomly propose one of the four possible moves, with probabilities:

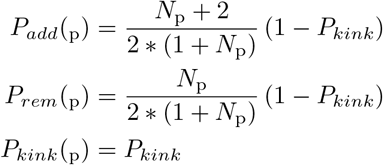

These are set to satisfy the detailed balance condition, that *P*_*add*_(*N*_*p*_) = *P*_*rem*_(*N*_*p*_ + 2) and *P*_*rem*_(*N*_*p*_) = *P*_*add*_(*N*_*p*_ *−* 2); since *P*_*kink*_ does not change the length of the polymer it can be held constant. After proposing a move we check for self-avoidance, and that the polymer meets the length constraint *N*_*p*_ < *N*_*max*_, if either of these are not satisfied we reject the move. If the move is not rejected, we compute the total energy from H_*poly*_ and accept with the Metropolis criteria 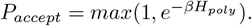.

#### Simulation of bulk system

Simulation of our bulk system closely follows the bulk system used in [15]. We simulate a two-component fluid of short polymers whose length *N*_*b*_ = 20 monomer units in all simulations. We update the position of a single bulk molecule via kink moves, which are analogous to those we propose in the long polymer. Bulk molecules also perform reptation moves where a monomer is removed from one end of the molecule and placed in a position adjacent to the monomer at the other end. These moves conserve length and are proposed randomly with uniform probability. Our bulk is typically held at fixed chemical potential. To this end we separately simulate a reservoir of non-interacting molecules and exchange particles with this reservoir with rate 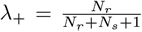 and 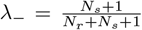 to ensure detailed balance. Particle exchanges and moves are accepted with the Metropolis probability given by the bulk Hamiltonian.

#### Multi-component polymers

Our simulations of polymers with multiple monomer types use the same move-set as the single-monomer simulations. We first equilibrate a single-monomer polymer for several million MCS, then impose monomer sequence on the polymer. We track the number of monomers per ‘segment’, and moves are allowed to change the monomers-per-segment up to *±*Δ: i.e if we set a segment of 50 monomers of the same type, and Δ = 10, this segment must have 40 *−−*60 monomers. We set Δ = 12 in all simulations displayed in Figure 4. Any move that brings the segment outside of this window is rejected. Otherwise the simulations are identical up the Hamiltonian, which now reflects the differences in binding to the bulk

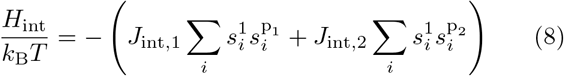

where, for example, *J*_int,1_ denotes interactions of the first monomer type with the bulk, and 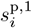 denotes the occupancy of the i^th^ lattice site for monomers of type 1.

